# Enhancing ecosystem service provision through the silvicultural management of European black pine stands from afforestation and reforestation

**DOI:** 10.1101/2025.01.31.635893

**Authors:** Elia Vangi, Sandro Sacchelli, Susanna Nocentini, Manuela Plutino, Daniela Dalmonech, Alessio Collalti, Davide Travaglini, Piermaria Corona

**Author notes:** Corresponding author: Elia Vangi. geoLAB - Laboratory of Forest Geomatics, Dept. of Agriculture, Food, Environment and Forestry, Università degli Studi di Firenze, Via San Bonaventura 13, 50145, Firenze, Italy.

## Abstract

Afforestation and reforestation are integral components of the wider field of land management. When these initiatives integrate the diverse eco-biological, landscape, cultural, and socioeconomic characteristics of the intervention area they can achieve substantial environmental improvements also by improving ecosystem functions, commonly referred to as ecosystem services (ES). European black pines are some of most frequently used tree species for afforestation and reforestation in Mediterranean regions, thanks to their ability to grow in poor soil conditions and their resistance to environmental stressors. In this study, we adopted a validated process-based modeling approach to explore the effects of thinning intensity and frequency on the provision of some ES from European black pine stands, which have been established through afforestation and reforestation in Italy. We found a net financial gain when basal area removal reaches 25% with a 25-year thinning interval, highlighting the higher financial efficiency of more intensive interventions. Non-provisioning ES (erosion protection, carbon sequestration, and aesthetic/recreational value) tend to decrease with increased basal area removal and benefit from longer intervals between thinning. Remarkably, the economic values of aesthetic appeal and carbon sequestration far exceed those of timber production and erosion protection, regardless of thinning regime. Based on our results, we claim that strategic, long-term planning of thinning operations is essential to ensure a balanced trade-off between wood production and other ES while maintaining the cost-effectiveness of operations. Ultimately, our approach can provide guidelines for forest managers to ensure the provision of multiple ES.

## 1. Introduction

Afforestation and reforestation fall within the broader scope of land management. For these practices to be effective, planting design must consider the natural processes that guide the spatial establishment of forest vegetation. This involves identifying the most appropriate intervention strategies for varying operational contexts and ensuring alignment with environmental, socioeconomic, and cultural objectives and conditions. Approaches can range from fostering natural colonization (promoting the development of biologically stable and self-sustaining ecosystems) to applying silvicultural techniques to establish semi-natural forests in the long term or even using intensive forest management methods typical of commercial forestry (Puettmann et al., 2015).

When afforestation and reforestation initiatives integrate the diverse eco-biological, landscape, cultural, and socio-economic characteristics of the intervention area during the planning, design, and establishment phases, they can achieve substantial environmental improvements by fostering the creation of semi-natural forests (Vacek et al., 2023). In Europe, and especially in Mediterranean regions, Austrian pine (*Pinus nigra* Arnold) and Corsican pine (*Pinus laricio* (Poir.) Maire), collectively known under the label of European black pines, are some of the most frequently used tree species for afforestation and reforestation. They cover 4.4% of the European forest area, and they are some of the most planted species outside their native range. European black pine plantations are mainly targeted to improve degraded landscapes. Thanks to their ability to grow in poor soil conditions and their resistance to environmental stressors such as drought and heat waves, these species are highly suitable in different habitats and resilient to climate change (Vacek et al., 2023). In Italy, these plantations, largely established after World War II, now span over 120,000 hectares nationwide, according to the Italian National Forest Inventory (Gasparini et al., 2022).

Very few studies have examined the impact of silvicultural practices, particularly thinning operations, on the ecosystem functions, commonly referred to as ecosystem services (ES), provided by conifer stands originating from afforestation and reforestation projects targeted to improve degraded landscapes. Thinning is a silvicultural treatment that reduces tree density to enhance the vigor of the remaining trees and improves the economic viability of the stands. However, thinning operations tend to have relatively high costs in conifer stands established from afforestation and reforestation. On the other hand, thinning can increase the provision of various ES in pine plantations, as noted by Simon and Ameztegui (2023).

ES are increasingly influential in forest management decision-making and planning: they represent the benefits ecosystems provide to human society, sustaining life on Earth and supporting human needs (Corona and Alivernini, 2024). However, integrating ES into forest management is challenging due to the trade-offs and synergies that often exist among different ES (e.g., Croitoru, 2007; Duncker et al., 2012; Wolfslehner et al., 2019; Nocentini et al., 2022). The characteristics of forest stands and management decisions can significantly influence the provision of ES (e.g., Tomao et al., 2017; Corona et al., 2018; Simon and Ameztegui, 2023), for example: thinning treatments, which modify the composition and horizontal and vertical structure of forest stands, affect the stand’s productive and protective functions and increase fire resistance; by reducing competition among trees, thinning can influence biodiversity, water cycle components, and recreational and landscape functions; it also impacts elemental cycles, such as carbon assimilation and nutrient mineralization, as solar radiation distribution on the forest floor changes, and the hydrological cycle (Saponaro et al., 2025).

In this study, we adopted a robust modeling approach to explore the effects of thinning intensity and frequency (from now on also referred to as thinning regimes) on the provision of ES in European black pine stands established through afforestation and reforestation. To achieve this, we used a process-based forest model, a key tool for analyzing the impacts of silvicultural management strategies on stand dynamics (Collalti et al., 2018; Dalmonech et al., 2022; Mahnken et al., 2022; Vangi et al., 2024a,b). Specifically, we implemented various thinning regimes using the 3D-CMCC-FEM v 5.6 model (Collalti et al., 2018; Testolin et al., 2023). Subsequently, we assessed the provision of multiple ES, including wood production, erosion control, carbon sequestration, aesthetic/recreational value and prevented structural damage, and their monetary values through the integration of economic models. The monetary valuation of the selected ES was based on environmental appraisal methods and on the concept of social (non-market) utility value (Pearce et al., 2003).

This integrated approach to evaluating ES provision under different thinning regimes offers valuable insights. The results can inform the silvicultural management of European black pine plantations, supporting their long-term sustainability and multifunctional role in delivering ES.

## 2. Materials and methods

### 2.1. Study area

The analyses were conducted in three study areas featuring European black pine plantations in Italy: Monte Amiata and Rincine in the Tuscany region, and Varco San Mauro in the Calabria region (Figure 1).

**Figure 1.**
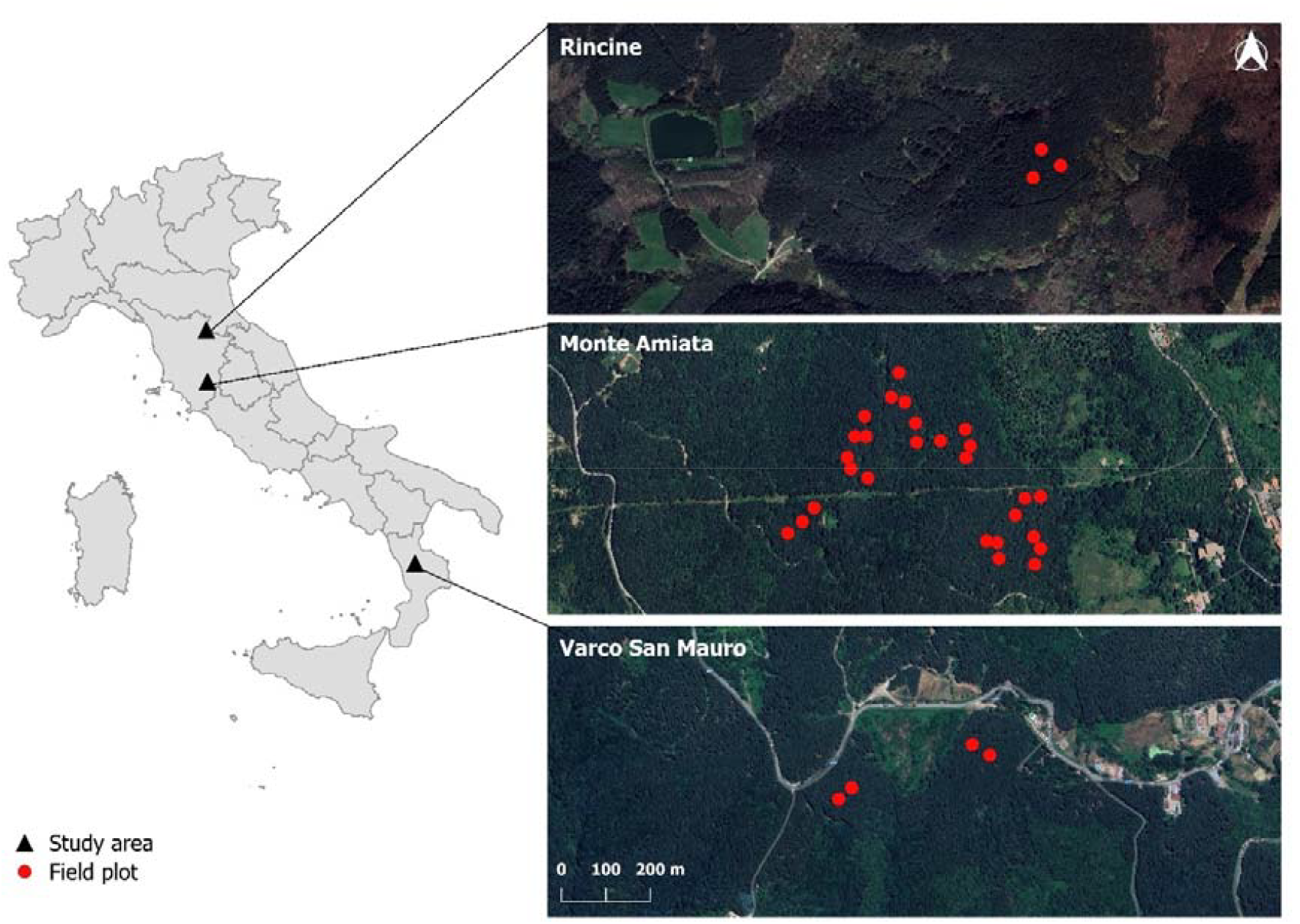
Location of the study areas and distribution of experimental field plots.

The Monte Amiata plantation is located on a southwest-facing aspect at an altitude of approximately 750 m a.s.l., with an average slope of 15%. In 2016, 27 circular plots, each with a radius of 15 m, were identified. For each tree within the plots, height and diameter at breast height were measured, and the growing stock volume was calculated using species-specific allometric models. These measurements were taken before and after applying three experimental management protocols: thinning from below by removing mostly dominated trees, selective thinning by selecting candidate trees and removing their competitors’ trees at the crown level, and a control (no treatment). All plots were remeasured in 2023, when the stand reached 52 years of age.

The Rincine plantation is situated at an altitude of approximately 1050 m a.s.l., with a southwest-facing aspect and an average slope of 70%. In 2008, three experimental plots, each measuring 50×50 m, were established and measured using the same protocol as at Monte Amiata. These measurements were conducted both before and after implementing three experimental silvicultural treatments: thinning from below, fellings carried by opening very small gaps to foster natural regeneration, and a control (no treatment). The stand was remeasured in 2024 when it was 51 years old.

The Varco San Mauro plantation is located within the Sila Plateau Forest at an altitude of 1118 m a.s.l. on south and southwest-facing slopes. The experimental protocol involved establishing four plots, each measuring 30×30 m, where two treatments were applied: selective thinning and a control (no treatment). The same measurement protocol as in the other two sites was used. Measurements were conducted in 2007, both before and after thinning, with a final survey performed in 2023, when the stand reached 63 years of age.

### 2.2. 3D-CMCC-FEM Model

The ‘*Three-Dimensional - Coupled Model Carbon Cycle - Forest Ecosystem Module*’ (3D-CMCC-FEM v 5.6) is a process–based model that simulates the biochemical and biophysical processes at ecosystem level, and structural dynamics of forest stands on time scales ranging from daily to decadal depending on the process to simulate. It is designed to model carbon, nitrogen, and water cycles in forest ecosystems and can simulate silvicultural interventions in pure or mixed even-aged and uneven-aged stands, including those with complex (multi-layered) structures, under present-day climate and climate change scenarios (Dalmonech et al., 2022; Vangi et al., 2024a, b; Collalti et al., 2016, 2018).

Photosynthesis is modeled using the biogeochemical approach developed by Farquhar, von Caemmerer, and Berry (Farquhar et al., 1980), with separate calculations for light and shade leaves (de Pury and Farquhar, 1997). The model accounts for the acclimation of leaf photosynthesis to temperature increases and simulates autotrophic respiration (RA) by distinguishing between the maintenance costs of existing tissues and the synthesis costs of new tissues. Maintenance respiration is controlled by the nitrogen content in living tissues (a stochiometrically fixed fraction of carbon content) and temperature.

Net Primary Productivity (NPP) is calculated as the difference between Gross Primary Productivity (GPP) and RA. Annual NPP is allocated to various compartments, including biomass production and the non-structural carbon pool (NSC), which stores starch and sugars for use during periods of negative carbon balance (when RA>GPP). Summer values of NSC below a set threshold, leads to progressive crown defoliation. If NSC reserves are depleted and not replenished, the model predicts tree mortality based on McDowell et al.’s (2008) carbon starvation hypothesis. The model incorporates species-specific phenological and allometric patterns as a function of stand age and (and because of) biomass accumulation. It is initialized using structural data from the forest stand (e.g., mean stem diameter, mean tree height, stand age, and stand density) and is driven by daily climate variables (e.g., temperature, precipitation, solar radiation, relative humidity) and annual atmospheric CO_2_ concentration values (μmol mol^−1^). Other input data include soil texture, soil depth, and site elevation. More details are reported in Collalti et al. (2024).

The forest ecosystem model has been tested and evaluated over several sites in Italy and Europe showing good performances in simulating carbon fluxes and structural variables (Collalti et al., 2016; Dalmonech et al., 2022; 2024; Testolin et al., 2023; Morichetti et al., 2024), and when compared to other forest models (Mahnken et al., 2022; Saponaro et al., 2025).

### 2.3. Climate data

The 3D-CMCC-FEM model requires daily climate data for the simulation period. For this study, the model was driven by daily climate data derived from the ERA5 reanalysis, downscaled specifically for Italy (Raffa et al., 2023). Downscaling was performed using the regional climate model COSMO5.0_CLM9 with INT2LM 2.06 (Rockel et al., 2008), which enhanced the spatial resolution from 31×31 km to 2.2×2.2 km and retained the original hourly timescale of ERA5 data.

The meteorological variables required for model simulation included daily minimum, maximum temperatures (Tmin, Tmax in °C), total daily precipitation (Pr in mm h□^1^), daily mean net surface shortwave radiation (Rg in MJ m□^2^ h□^1^), and the relative humidity (RH, in %). From the average temperature (Tav) and Td (the dew point), the daily relative humidity was calculated using the R Humidity package (Cai, 2019). Hourly data were aggregated to a daily timescale by calculating the mean values, except for precipitation, for which the daily sum was computed.

All climate data required for the model were extracted for each field plot for the period 1981-2023, which, despite the low climate change signal already observable in this time span, can be considered the current climate. Global annual values of atmospheric CO_2_ concentration were extracted from the PROFOUND database (Reyer et al., 2020) for the same period (1981-2023).

To simulate stand growth without the influence of future climate change and thus emphasize the effect of the different thinning regimes, a long climate record was created by detrending the downscaled historical climate data from 1981-2023 and repeated cyclically from 1981 up to the year 2100. The same procedure was applied to atmospheric CO_2_ concentration records.

### 2.4. Soil Data

Soil depth and texture (percentages of clay, silt, and sand) were obtained from the national soil database developed by the Soil Cartography Laboratory of the Italian Council for Agricultural Research and Agricultural Economics (Costantini and Dazzi, 2013; Corona et al., 2023). This database, with a spatial resolution of 250 m, consists of four layers representing soil depth (in cm) and the percentages of clay, silt, and sand within the first meter of depth.

### 2.5. Model simulation scenarios and evaluation

Firstly, to ensure that the model accurately reproduced the growth trends of the investigated stands, including the response to thinning from below, the 3D-CMCC-FEM was validated using the downscaled climate data and field measurements collected during two consecutive survey campaigns (pre- and post-intervention). The simulations were initialized with data from the first survey campaign (pre-intervention) conducted in the control plots (12 plots: nine in Monte Amiata, two in Varco San Mauro, and one in Rincine), where no silvicultural interventions had been applied, and in the thinned from below plots (10 plots: nine in Monte Amiata and one in Rincine). The data needed for initializing the simulations were species, age and structural data such as mean tree diameter at breast height (DBH), mean tree height (HEIGHT) and the number of trees in the stand (TREE). A comprehensive report on the utilized data can be found in Corona et al. (2025). All the simulations were stopped at the year of the second measurement. The simulation outputs for the year of the second measurement were then compared with the corresponding field-observed values for key variables: Figures 2 and 3 illustrate the strong agreement between the predicted and observed values, confirming the model’s reliability and support its use in this study, both for no management and thinning scenarios.

**Figure 2.**
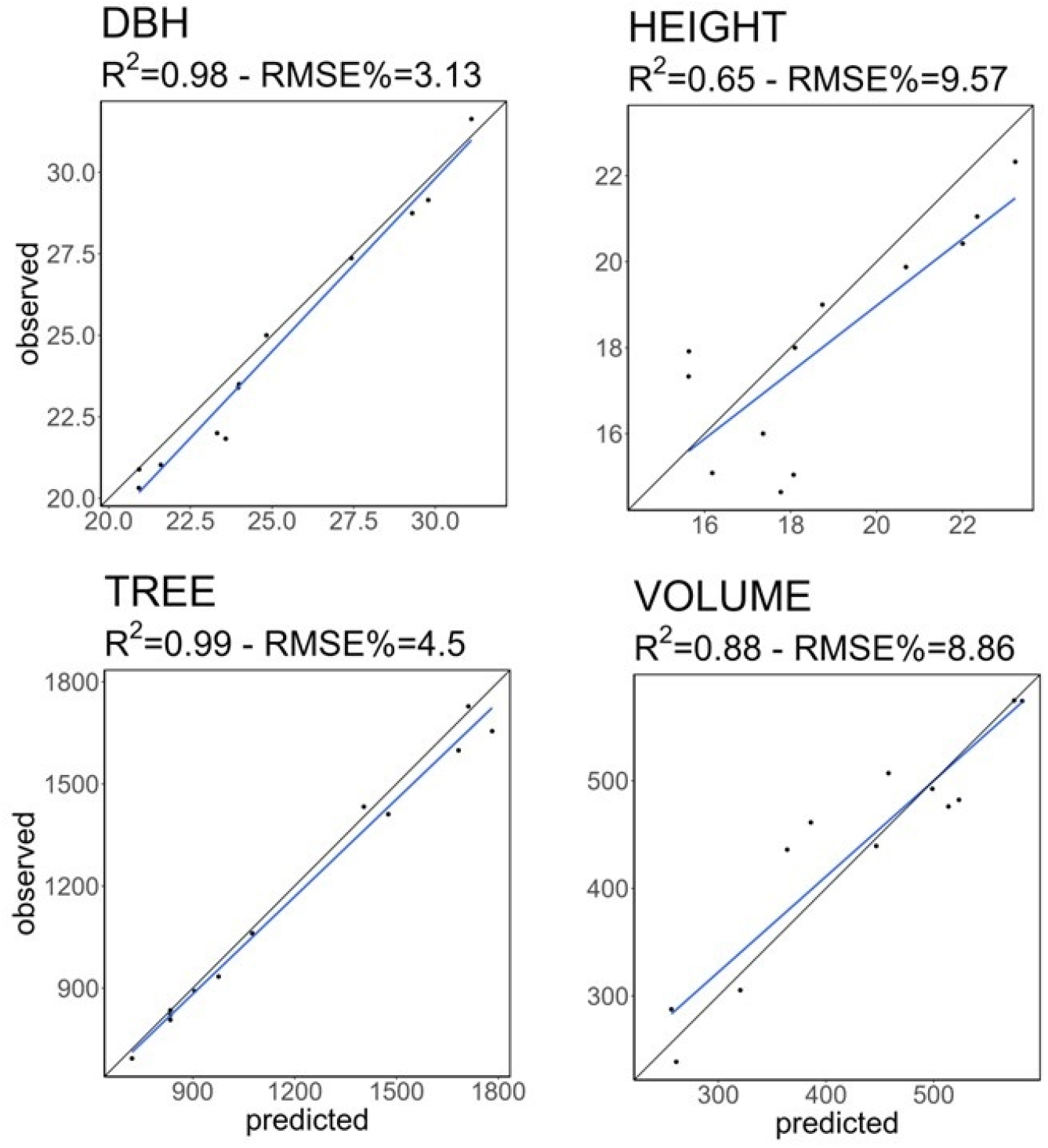
Values predicted by the 3D-CMCC-FEM model (x-axis) vs. observed values (y-axis) for selected variables (DBH = mean tree diameter at breast height; HEIGHT = mean tree height; TREE = number of trees; VOLUME = wood volume) in the control plots.

**Figure 3.**
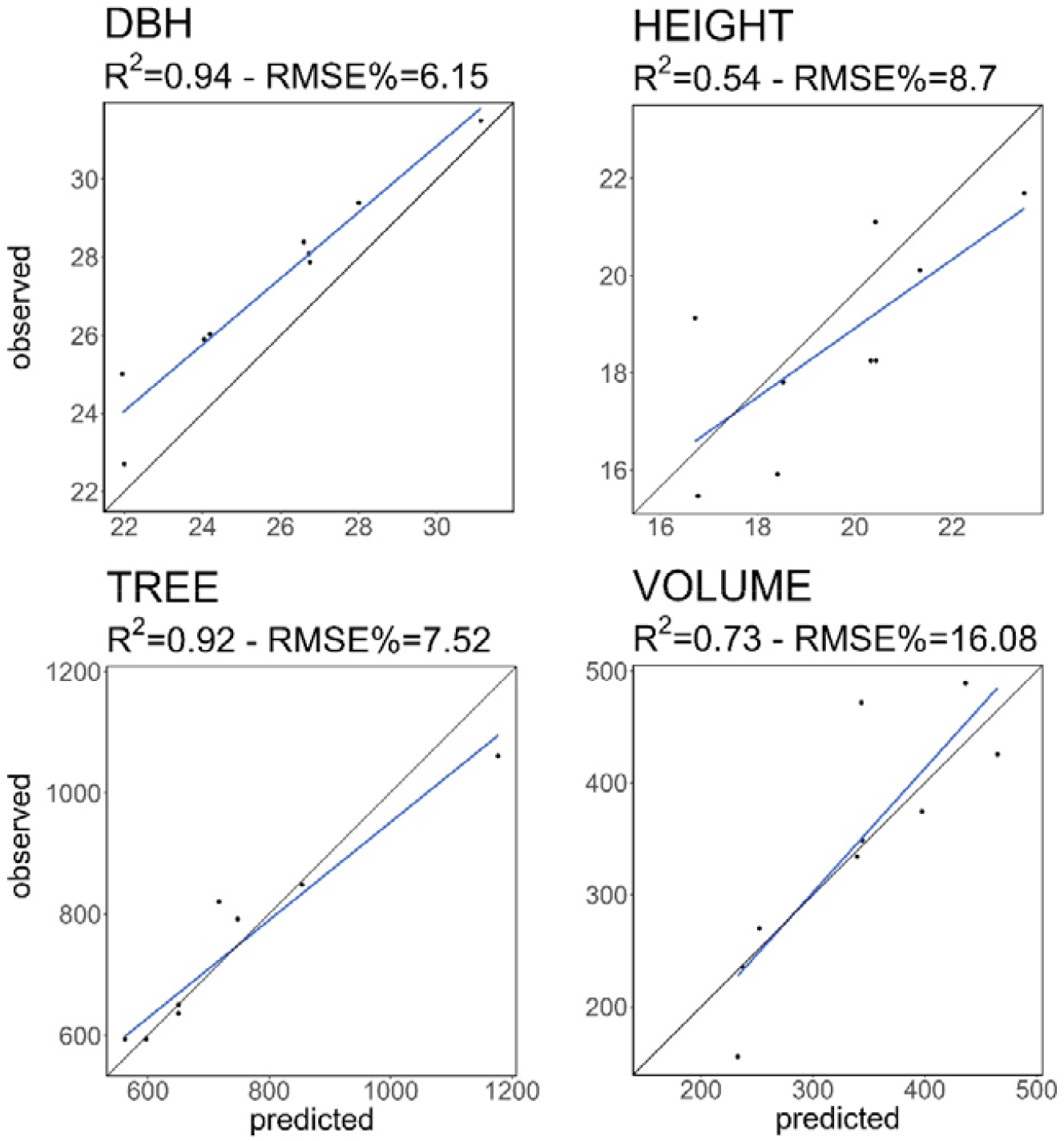
Values predicted by the 3D-CMCC-FEM model (x-axis) vs. observed values (y-axis) for selected variables (DBH = mean tree diameter at breast height; HEIGHT = mean tree height; TREE = number of trees; VOLUME = wood volume) in the thinned from below plots.

To evaluate stand dynamics of the 34 plots (27 at Monte Amiata, three at Rincine, four at Varco San Mauro) and the associated provision and economic quantification of ES in European black pine plantations, 20 thinning from below regimes were simulated using data collected from the experimental plots. These regimes were distinguished by thinning intensity (percentage of stand basal area to remove based on total stand basal area) and frequency (years between interventions), with combinations of thinning frequency ranging from 10 to 25 years (in 5-year increments) and thinning intensity varying from 15% to 35% of basal area (in 5% increments). In all thinning regimes, thinning from below was simulated by removing trees from smallest to largest until the target thinning intensity was reached. An additional non-intervention scenario was included as a control, resulting in a total of 20 thinning regimes (4 frequencies × 5 intensities) + 1 control. The focus of this assessment was to analyze the sole impact of thinning interventions on the selected ES, described in par 2.6. To isolate this effect, the rotation period was set such that final harvesting was excluded from the simulation period.

For the long-term simulations, each plot was initialized using the structural variable values measured in the pre-intervention survey. Simulations were conducted using the R3DFEM R package (Vangi et al., submitted), which provides an R wrapper for the 3D-CMCC-FEM model. These simulations covered the period from the year of the first survey at each study site to the year 2100, corresponding to a time span of approximately 83-93 years, depending on the site. The key variables selected for the economic quantification of ES included: DBH, measured in centimeters (cm); HEIGHT, measured in meters (m); stand basal area (B), measured in square meters per hectare (m^2^ ha□^1^); above-ground carbon content (AGC), measured in mega grams of carbon per hectare (MgC ha□^1^); canopy cover (C), expressed as a unitless proportion; harvested wood volume, measured in cubic meters per hectare per year (m^3^ ha□^1^ y□^1^).

### 2.6. Economic quantification of ES

The model outputs were used to estimate the provision of ES and quantify their economic value. The economic efficiency of thinning interventions was evaluated for each year of the simulation period. To underscore the role of thinning in enhancing stand stability, an assessment of avoided economic damage was conducted by comparing the outcomes with those of the non-intervention (control) scenario.

#### 2.6.1. Wood production

The financial value of wood production is quantified by calculating the stumpage value (SV) of harvested wood volume in year n. The SV is a typical transformation value that is defined, by the classic forestry economic-appraisal approaches, as the value of standing trees, or the difference between revenues (I) and costs (S) of the production process:

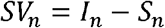

Revenues quantification considers both obtainable volume and average selling price of wooded assortments. Cost calculation involved an analysis of the regional price lists (Tuscany and Calabria) for the reference operations. Specifically, thinning operations in coniferous forests were used, including cutting, debranching, extraction, clearing and arrangement of the site as well as general expenses (direction cost, administrative cost, interests; Bernetti and Romano, 2007).

The Net Present Value for provision service (NP_SV) is calculated with the formula:

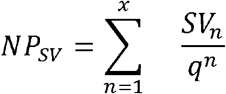

where *q* =1+ *r*, with *r* equal to the interest rate (assumed as 2.5% in this study; Sartori et al., 2014).

#### 2.6.2. Aesthetic value

The aesthetic value of an area is closely tied to its ability to meet the recreational needs of the population. Drawing on a synthesis of extensive scientific literature on this topic, the calculation of aesthetic value is based on the work of Ribe (2009). In this study, various types of coniferous stands (mature, aged, and those subject to logging) are analyzed to determine their scenic value, as perceived by a sample of respondents, using the Ratio Scenic Beauty Estimate (RSBE). The RSBE is calculated as a function of the basal area per hectare (B) through the following polynomial model:

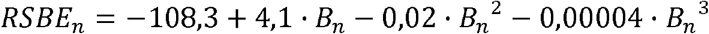

where *RSBE*_*n*_ indicates the value of RSBE in year *n*.

The RSBE values observed in Ribe (2009) vary approximately in the range +150 ÷ –150. This dimensionless value must therefore be managed to achieve the economic value of the aesthetic function. Benefit Transfer (BT) approach (Desvouges et al., 1998) is useful to quickly transfer the results from other case studies to the areas of interest. A meta-analysis based on Contingent Valuation, Discrete Choice Experiments and the Travel Cost Method were used, highlighting a Willingness to Pay (WTP) of 7.79 €/visit y^-1^ for recreational utility in coniferous forest for Italian context (Grilli et al., 2014). Reported WTP (7.79 €/visit y^-1^) can be cautiously considered as expressive of the maximum WTP for forests with optimal aesthetic value (RSBE = +150). With this hypothesis the annual WTP is quantified weighing the potential maximum WTP on the normalized RSBE (norm_RSBE_). The normalization occurred through the multicriteria approach of Compromise Programming (Romero and Rehman, 2003):

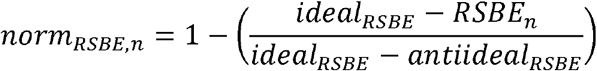

where *ideal*_*RSBE*_ and *antiideal*_*RSBE*_ represent, respectively, the ideal (+150) and non-ideal (–150) RSBE values as per Ribe (2009).

The aesthetic value (AV_*n*_) (€ ha^-1^ year^-1^) is then calculated as:

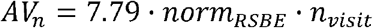

where *n_visit* is the estimated annual number of visits for the area under consideration. Given the absence of specific data in the various study areas, the value of *n_visit* was arbitrarily set to 150 for each examined stand.

The NPV for the aesthetic function (NP_AV) is quantified as follows:

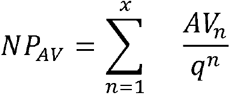

#### 2.6.3. Protection from erosion

The model is based on the quantification of avoided soil erosion due to forest cover against the effect of atmospheric precipitation. The economic value of the avoided erosion is then derived from the correlation with the price of sediment removal from potential basins located downstream of the forest area (Sacchelli et al., 2021).

The amount of erosion was computed through the modified version of the Revised Universal Soil Loss Equation (RUSLE2015) (Panagos et al., 2015a). The RUSLE2015 formula estimates soil loss (E, expressed in t ha^-1^ year^-1^), applying five input factors: rainfall erosivity (R), soil erodibility (K), forest canopy cover (C), topography factor (LS) and support practices (P):

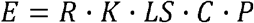

The geodata applied in the equation are freely available upon request in the European Soil Data Center (ESDAC) (https://esdac.jrc.ec.europa.eu/resource-type/soil-threats-data) (Panagos et al., 2022).

In this work, R, K and LS factors were kept constant given their very low variation in medium-long term and their low influence in the forest rotation. The P factor is not included due to fragmentary applications in the forestry sector (Panagos et al., 2015a).

The assessment of avoided erosion in year *n* is based on the difference between the factor *C*_*n*_ with forest and the factor *C*_*0*_ in the hypothesis of absence of forest cover (θ*n*=0). The value of *C* is quantified according to the formula reported in Panagos et al. (2015b):

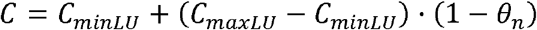

with *C*_*minLU*_ and *C*_*maxLU*_, respectively, minimum and maximum values of C for forests (Panagos et al., 2015b) and θ_*n*_ fraction of canopy cover at ground level in year *n*.

The avoided erosion in year *n* is then calculated as:

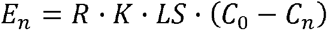

*E*_*n*_ was recalibrated with the application of the Sediment Delivery Ratio (SDR) (De Rosa et al., 2016) a coefficient allowing for quantification of actual soil debris from the basin slope to reservoir:

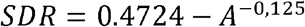

with *A* corresponding to the size of the catchment area in km^2^.

The monetary value of protection from erosion (PV) is based on the unitary cost (α) of sediment removal from artificial basins or reservoirs as established in Palmieri et al. (2014) (29.29 € t^-1^):

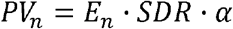

The NPV of the protective function (NP_PV) can be finally calculated as:

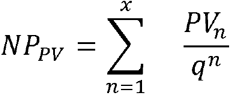

#### 2.6.4. Carbon storage

The estimate of carbon value starts from the growing stock volume and, with allometric equations and Biomass Expansion Factors (BEFs), led to the calculation of the above ground biomass (AGB, expressed in m^3^ ha^-1^) (Vitullo et al., 2007; Vangi et al., 2023). The total carbon is obtained by multiplying the biomass by its carbon content (0.47 grams of C per gram of dry matter) (MgC ha^-1^). To obtain the mass of stored CO_2_, the mass of carbon is multiplied by the β coefficient of 3.67 (Federici et al., 2008).

The European Emission Trading System (ETS) market was applied to derive carbon trading price (γ) (https://tradingeconomics.com/commodity/carbon).

The monetary annual value of the carbon storage function (CV) is given by (Sacchelli, 2018):

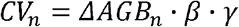

where ΔAGB represents the variation of AGB from year n-1 to year n; the CV_n_ equation precautionary hypothesises that – for the harvesting year – the carbon contained in the removed biomass is deducted from the quantification, without knowing the final use of the woody assortments.

The NPV of the externality (NP_CV) is obtained as:

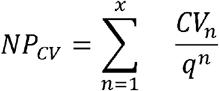

#### 2.6.5 Total Economic Value

The Net Present Total Economic Value (NP*_*TEV) is derived from the sum of the discounted value of the four ecosystem utilities:

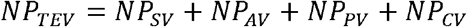

#### 2.6.6. Analysis of avoided damage

Thinning can provide significant benefits to the structural stability of stands (Hanewinkel et al., 2013; Suliman and Ledermann, 2025). Stability improvements can be quantified in both biophysical and economic terms. Within this context, assessing the potential indirect effects of thinning, such as avoided damage, is particularly relevant. Avoided damage can be evaluated by comparing the probability of adverse effects in stands without interventions (control) to those with interventions.

One of the key parameters for maintaining the structural stability of coniferous stands is the mean tree height – to - mean tree diameter at breast height ratio (H/D, see Slodicak and Novak, 2006). Building on the approach of Mickovski et al. (2005) and focusing specifically on wind damage, a risk trend associated with the H/D ratio can be identified. Notably, Mickovski et al. (2005) define five risk classes based on H/D ratio thresholds, ranging from <70 to >90. Starting from this categorization, the first step to calculate the missed damages was the quantification of the average value of the H/D ratio for each thinning regime; subsequently, with the aim of identifying a stability coefficient of the stands (*δ*), the H/D ratio was normalized in the range 0-1 with the compromise programming technique (Romero and Rehman, 2003). The ideal value was set equal to 50, while the anti-ideal value was equal, for each experimental plot, to the average H/D ratio of the non-intervention scenario. Finally, the avoided damage results from the combination (joint probability), for each plot and thinning regime, between the total economic value, the probability of stability increase linked to thinning, and the probability of extreme winds (λ). The λ coefficient represents the annual probability of winds potentially causing damage to the stands, related to the reference period (*µ*, years); the value was extracted at cartographic level for each plot with zonal statistics operations starting from the geodata of the research by Sacchelli et al. (2018).

The expected avoided damage *E(AD)* in the case of thinning is therefore quantified as: where *δ*_*no_thin*_ and *δ*_*thin*_ are, respectively, the stability coefficient of the stands (normalized H/D ratio) for the j-th scenario without and with intervention.

## 3. Results

### 3.1. Stand dynamics

The simulation outputs for varying thinning regimes applied to the 34 experimental plots of European black pine across the study areas are presented below. Figures 3 and 4 illustrate the trends in DBH and AGC over the simulation period for each combination of thinning frequency and intensity at each site.

**Figure 4.**
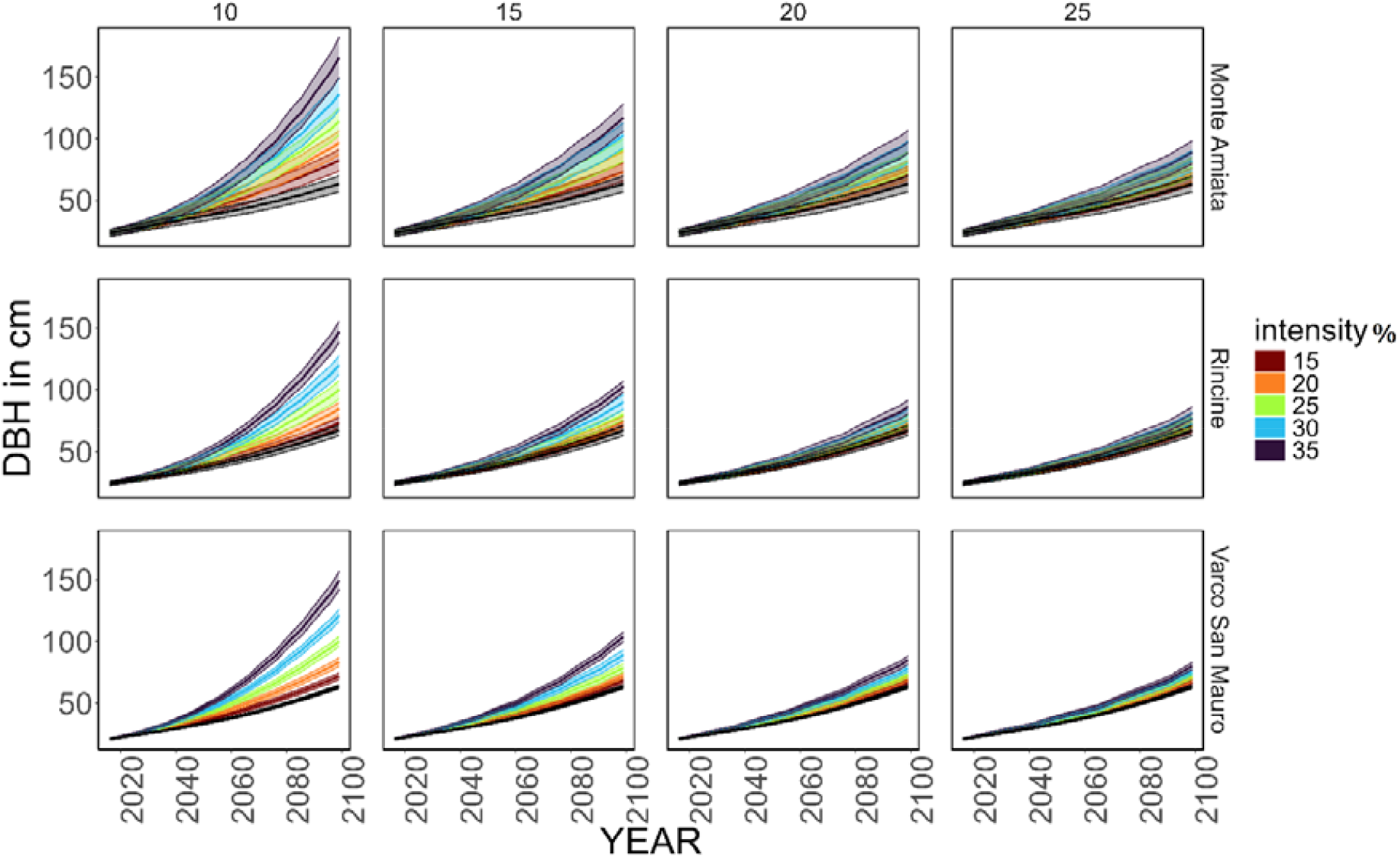
Average DBH (in cm) time series for each combination of site and thinning frequency (every 10, 15, 20, and 25 years), computed from the DBH of all simulated plots. Colors map different thinning intensities (removal of 15, 20, 25, 30, and 35% of basal area); the grey color is for the non-intervention (control) scenario. The shaded area represents one standard deviation from the mean value of all simulated plots.

The greatest increments in DBH and HEIGHT by the end of the simulation period (2100) were observed across all sites with a thinning frequency of 10 years at 35% intensity. Specifically, DBH growth reached 142 cm, 127 cm, and 122 cm, while HEIGHT growth reached 24 m, 25 m, and 23 m for Monte Amiata, Varco San Mauro, and Rincine, respectively. These were followed by a thinning frequency of 10 years at 30% intensity.

In contrast, AGC and basal area exhibited greater growth under a longer thinning frequency of 25 years, combined with the same 35% intensity (AGC increase of 73, 52, and 30 MgC ha□^1^ for Monte Amiata, Varco San Mauro, and Rincine, respectively). Overall, heavy-intensity thinning regimes applied at short intervals yielded the highest DBH increments, with values doubling those observed under the no-management scenario. However, these regimes also resulted in substantial wood removal, leading to lower basal area and AGC values. Consequently, the highest volume of harvested woody products across all thinning regimes was achieved with a 10-year frequency and 35% intensity, exceeding 2100, 1980 and 1680 m^3^ ha□^1^ over approximately 90 years for Monte Amiata, Varco San Mauro, and Rincine, respectively. Conversely, the lowest volume of harvested woody products (excluding the control scenario with no intervention) was recorded under the least intensive thinning regime (15%) combined with the longest frequency (25 years).

### 3.2 Economic quantification of ES

The results of the thinning regimes for the examined plantations indicate that NP_SV increases with both the amount of basal area removed and the length of the thinning interval (Tables 1, 2, and 3). Analyzing NP_SV in relation to thinning interval and wood removal, it was observed that, at the Monte Amiata site, NP_SV remained negative up to a removal threshold of 20% of the basal area. For higher levels of wood removal, a positive monetary value was achieved when the thinning interval was at least 15 years (Figure 6).

**Table 1.** Economic results for the plantation of Monte Amiata for each thinning regime (frequency_intensity) including the non-intervention (control) scenario.

| Thinning regime<br>(years_%) | NP_SV<br>(€ ha<sup>-1</sup>) | NP_AV<br>(€ ha<sup>-1</sup>) | NP_PV<br>(€ ha<sup>-1</sup>) | NP_CV<br>(€ ha<sup>-1</sup>) | NP_TEV<br>(€ ha<sup>-1</sup>) | Average<br>H/D ratio | Norm<br>H/D ratio | Damage<br>thinning<br>(€ ha<sup>-1</sup>) | Damage<br>no<br>thinning<br>(€ ha<sup>-1</sup>) | Avoided<br>damage<br>(€ ha<sup>-1</sup>) | %_avoided<br>dam TEV |
| --- | --- | --- | --- | --- | --- | --- | --- | --- | --- | --- | --- |
| 10_15 | -11250 | 25933 | 141 | 30671 | 45495 | 65 | 0.78 | 8347 | 10700 | 2353 | 5 |
| 10_20 | -7377 | 24622 | 142 | 27394 | 44781 | 63 | 0.66 | 6946 | 10532 | 3586 | 8 |
| 10_25 | -3164 | 23188 | 141 | 24353 | 44517 | 60 | 0.53 | 5541 | 10470 | 4930 | 11 |
| 10_30 | -625 | 21713 | 140 | 21472 | 42700 | 57 | 0.39 | 3961 | 10043 | 6082 | 14 |
| 10_35 | 1072 | 20143 | 139 | 18652 | 40007 | 55 | 0.25 | 2378 | 9410 | 7031 | 18 |
| 15_15 | -8224 | 26803 | 142 | 33762 | 52482 | 67 | 0.87 | 10742 | 12344 | 1601 | 3 |
| 15_20 | -5033 | 25936 | 142 | 31168 | 52213 | 65 | 0.79 | 9707 | 12280 | 2573 | 5 |
| 15_25 | -1305 | 24931 | 141 | 28665 | 52433 | 63 | 0.70 | 8673 | 12332 | 3659 | 7 |
| 15_30 | 1313 | 23824 | 140 | 26221 | 51498 | 61 | 0.60 | 7301 | 12112 | 4812 | 9 |
| 15_35 | 3674 | 22668 | 139 | 23842 | 50323 | 59 | 0.50 | 5917 | 11836 | 5919 | 12 |
| 20_15 | -6708 | 27125 | 142 | 34304 | 54863 | 67 | 0.91 | 11685 | 12904 | 1219 | 2 |
| 20_20 | -3795 | 26511 | 142 | 32463 | 55322 | 66 | 0.85 | 11092 | 13012 | 1919 | 3 |
| 20_25 | -1042 | 25755 | 141 | 30428 | 55281 | 65 | 0.79 | 10231 | 13002 | 2771 | 5 |
| 20_30 | 1375 | 24879 | 140 | 28122 | 54516 | 64 | 0.71 | 9124 | 12822 | 3698 | 7 |
| 20_35 | 3768 | 23972 | 140 | 26134 | 54014 | 62 | 0.63 | 8038 | 12704 | 4666 | 9 |
| 25_15 | -6402 | 27275 | 142 | 35190 | 56204 | 68 | 0.92 | 12183 | 13219 | 1037 | 2 |
| 25_20 | -4116 | 26768 | 142 | 33659 | 56452 | 67 | 0.88 | 11697 | 13278 | 1581 | 3 |
| 25_25 | -871 | 26156 | 141 | 31909 | 57336 | 66 | 0.83 | 11204 | 13485 | 2281 | 4 |
| 25_30 | 1476 | 25467 | 140 | 30131 | 57215 | 65 | 0.77 | 10412 | 13457 | 3045 | 5 |
| 25_35 | 3665 | 24679 | 140 | 28329 | 56813 | 63 | 0.71 | 9443 | 13362 | 3919 | 7 |
| Control | 0 | 28324 | 143 | 38142 | 66609 | 69 | 1.00 | 15666 | 15666 | 0 | 0 |

**Table 2.**
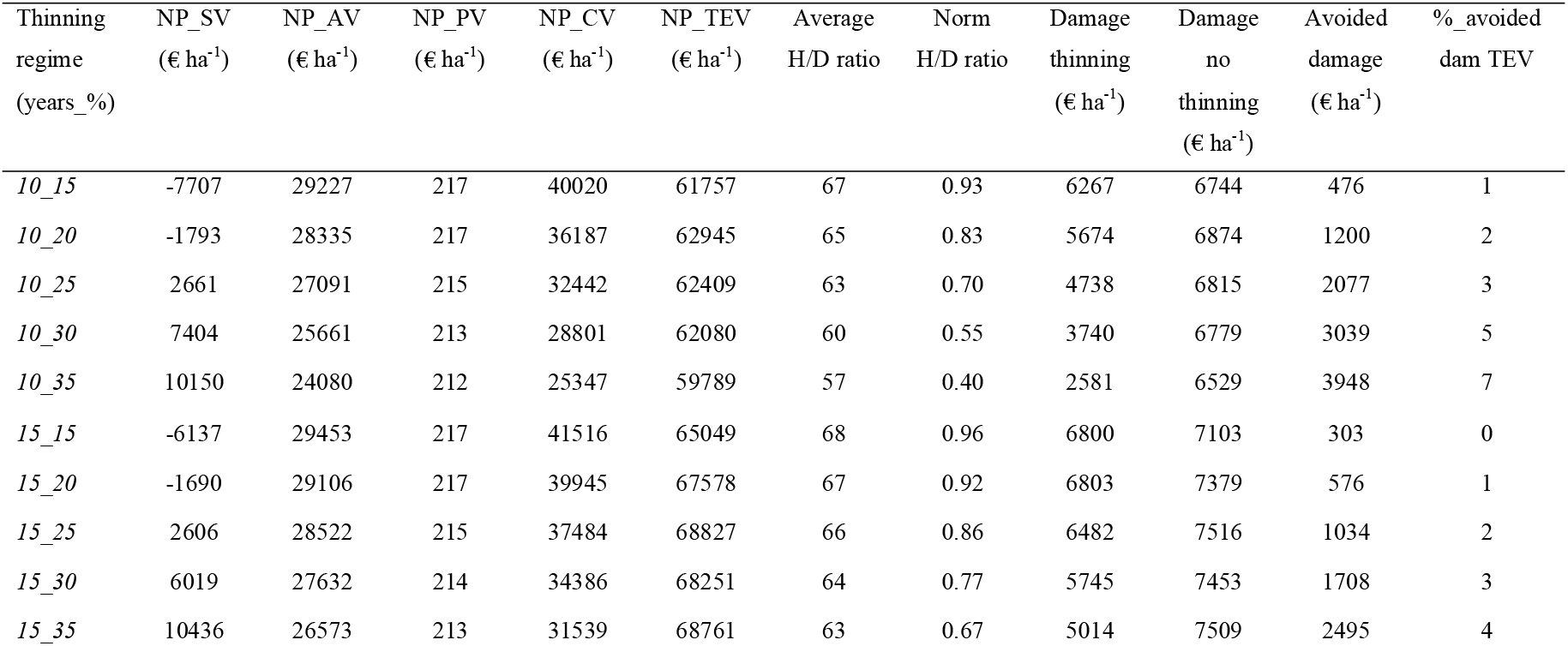
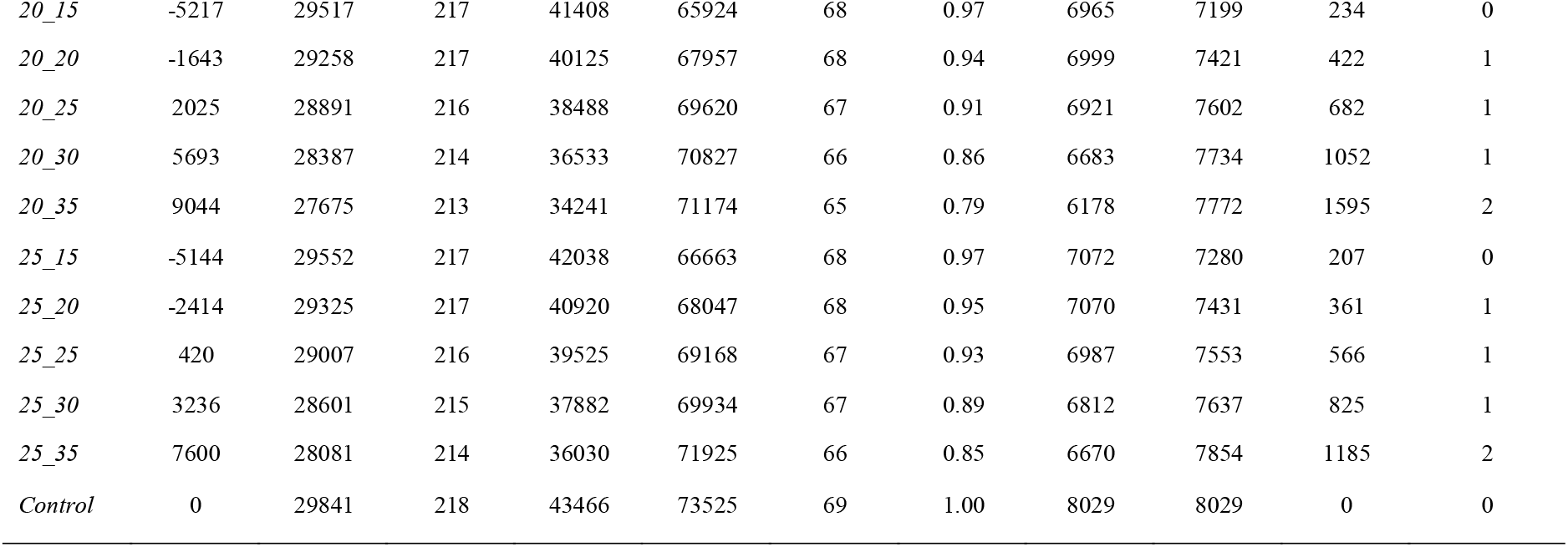
Economic results for the plantation of Rincine for each thinning regime (frequency_intensity) including the non-intervention (control) scenario.

| Thinning regime<br>(years_%) | NP_SV<br>(€ ha<sup>-1</sup>) | NP_AV<br>(€ ha<sup>-1</sup>) | NP_PV<br>(€ ha<sup>-1</sup>) | NP_CV<br>(€ ha<sup>-1</sup>) | NP_TEV<br>(€ ha<sup>-1</sup>) | Average<br>H/D ratio | Norm<br>H/D ratio | Damage<br>thinning<br>(€ ha<sup>-1</sup>) | Damage<br>no<br>thinning<br>(€ ha<sup>-1</sup>) | Avoided<br>damage<br>(€ ha<sup>-1</sup>) | %_avoided<br>dam TEV |
| --- | --- | --- | --- | --- | --- | --- | --- | --- | --- | --- | --- |
| 10_15 | -7707 | 29227 | 217 | 40020 | 61757 | 67 | 0.93 | 6267 | 6744 | 476 | 1 |
| 10_20 | -1793 | 28335 | 217 | 36187 | 62945 | 65 | 0.83 | 5674 | 6874 | 1200 | 2 |
| 10_25 | 2661 | 27091 | 215 | 32442 | 62409 | 63 | 0.70 | 4738 | 6815 | 2077 | 3 |
| 10_30 | 7404 | 25661 | 213 | 28801 | 62080 | 60 | 0.55 | 3740 | 6779 | 3039 | 5 |
| 10_35 | 10150 | 24080 | 212 | 25347 | 59789 | 57 | 0.40 | 2581 | 6529 | 3948 | 7 |
| 15_15 | -6137 | 29453 | 217 | 41516 | 65049 | 68 | 0.96 | 6800 | 7103 | 303 | 0 |
| 15_20 | -1690 | 29106 | 217 | 39945 | 67578 | 67 | 0.92 | 6803 | 7379 | 576 | 1 |
| 15_25 | 2606 | 28522 | 215 | 37484 | 68827 | 66 | 0.86 | 6482 | 7516 | 1034 | 2 |
| 15_30 | 6019 | 27632 | 214 | 34386 | 68251 | 64 | 0.77 | 5745 | 7453 | 1708 | 3 |
| 15_35 | 10436 | 26573 | 213 | 31539 | 68761 | 63 | 0.67 | 5014 | 7509 | 2495 | 4 |
| 20_15 | -5217 | 29517 | 217 | 41408 | 65924 | 68 | 0.97 | 6965 | 7199 | 234 | 0 |
| 20_20 | -1643 | 29258 | 217 | 40125 | 67957 | 68 | 0.94 | 6999 | 7421 | 422 | 1 |
| 20_25 | 2025 | 28891 | 216 | 38488 | 69620 | 67 | 0.91 | 6921 | 7602 | 682 | 1 |
| 20_30 | 5693 | 28387 | 214 | 36533 | 70827 | 66 | 0.86 | 6683 | 7734 | 1052 | 1 |
| 20_35 | 9044 | 27675 | 213 | 34241 | 71174 | 65 | 0.79 | 6178 | 7772 | 1595 | 2 |
| 25_15 | -5144 | 29552 | 217 | 42038 | 66663 | 68 | 0.97 | 7072 | 7280 | 207 | 0 |
| 25_20 | -2414 | 29325 | 217 | 40920 | 68047 | 68 | 0.95 | 7070 | 7431 | 361 | 1 |
| 25_25 | 420 | 29007 | 216 | 39525 | 69168 | 67 | 0.93 | 6987 | 7553 | 566 | 1 |
| 25_30 | 3236 | 28601 | 215 | 37882 | 69934 | 67 | 0.89 | 6812 | 7637 | 825 | 1 |
| 25_35 | 7600 | 28081 | 214 | 36030 | 71925 | 66 | 0.85 | 6670 | 7854 | 1185 | 2 |
| Control | 0 | 29841 | 218 | 43466 | 73525 | 69 | 1.00 | 8029 | 8029 | 0 | 0 |

**Table 3.** Economic results for the plantation of Varco San Mauro for each thinning regime (frequency_intensity) including the non-intervention (control) scenario.

| Thinning regime<br>(years_%) | NP_SV<br>(€ ha<sup>-1</sup>) | NP_AV<br>(€ ha<sup>-1</sup>) | NP_PV<br>(€ ha<sup>-1</sup>) | NP_CV<br>(€ ha<sup>-1</sup>) | NP_TEV<br>(€ ha<sup>-1</sup>) | Average<br>H/D ratio | Norm<br>H/D ratio | Damage<br>thinning<br>(€ ha<sup>-1</sup>) | Damage<br>no<br>thinning<br>(€ ha<sup>-1</sup>) | Avoided<br>damage<br>(€ ha<sup>-1</sup>) | %_avoided<br>dam<br>TEV |
| --- | --- | --- | --- | --- | --- | --- | --- | --- | --- | --- | --- |
| 10_15 | -9785 | 28594 | 164 | 37970 | 56943 | 68 | 0.91 | 12988 | 14350 | 1362 | 2 |
| 10_20 | -4871 | 27635 | 164 | 34433 | 57361 | 66 | 0.80 | 11541 | 14455 | 2914 | 5 |
| 10_25 | 82 | 26390 | 163 | 30979 | 57614 | 63 | 0.67 | 9709 | 14519 | 4810 | 8 |
| 10_30 | 3646 | 25032 | 161 | 27716 | 56555 | 61 | 0.53 | 7503 | 14252 | 6749 | 12 |
| 10_35 | 7730 | 23567 | 160 | 24602 | 56060 | 58 | 0.37 | 5281 | 14127 | 8846 | 16 |
| 15_15 | -8012 | 28938 | 164 | 39725 | 60815 | 69 | 0.95 | 14496 | 15325 | 829 | 1 |
| 15_20 | -3578 | 28482 | 164 | 38065 | 63133 | 68 | 0.90 | 14388 | 15910 | 1521 | 2 |
| 15_25 | 300 | 27805 | 163 | 35705 | 63974 | 67 | 0.84 | 13520 | 16121 | 2602 | 4 |
| 15_30 | 3559 | 26909 | 162 | 33006 | 63636 | 65 | 0.75 | 11991 | 16036 | 4045 | 6 |
| 15_35 | 7701 | 25828 | 161 | 30130 | 63820 | 63 | 0.64 | 10291 | 16083 | 5792 | 9 |
| 20_15 | -6799 | 29032 | 164 | 39761 | 62159 | 69 | 0.96 | 14995 | 15664 | 669 | 1 |
| 20_20 | -3110 | 28694 | 164 | 38353 | 64101 | 69 | 0.93 | 15017 | 16153 | 1137 | 2 |
| 20_25 | 161 | 28242 | 163 | 36670 | 65236 | 68 | 0.89 | 14669 | 16440 | 1771 | 3 |
| 20_30 | 3420 | 27654 | 162 | 34689 | 65926 | 67 | 0.84 | 13960 | 16613 | 2653 | 4 |
| 20_35 | 6596 | 26940 | 161 | 32668 | 66365 | 66 | 0.77 | 12930 | 16724 | 3794 | 6 |
| 25_15 | -6104 | 29086 | 164 | 40416 | 63562 | 69 | 0.96 | 15448 | 16018 | 570 | 1 |
| 25_20 | -3678 | 28786 | 164 | 39270 | 64543 | 69 | 0.94 | 15320 | 16265 | 944 | 1 |
| 25_25 | -1184 | 28407 | 163 | 37888 | 65274 | 68 | 0.91 | 15028 | 16449 | 1421 | 2 |
| 25_30 | 1358 | 27944 | 162 | 36258 | 65722 | 68 | 0.88 | 14533 | 16562 | 2029 | 3 |
| 25_35 | 3953 | 27381 | 162 | 34461 | 65956 | 67 | 0.83 | 13811 | 16621 | 2809 | 4 |
| Control | 0 | 29608 | 165 | 42351 | 72123 | 70 | 1.00 | 18175 | 18175 | 0 | 0 |

**Figure 5.**
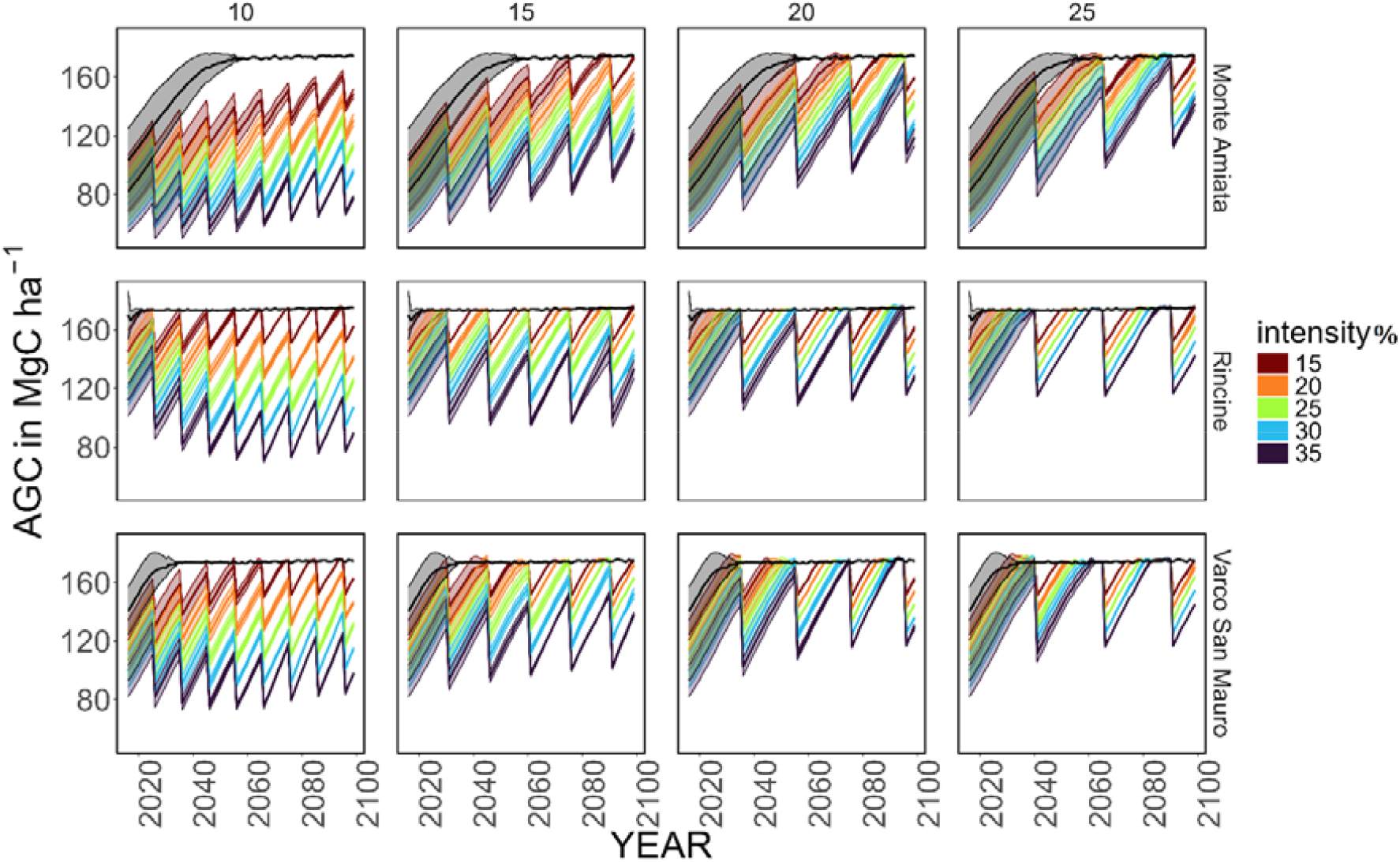
Average AGC (in MgC ha^-1^) time series for each combination of site and thinning frequency (every 10, 15, 20, and 25 years), computed from the AGC of all simulated plots. Colors map different thinning intensities (removal of 15, 20, 25, 30, and 35% of basal area); the grey color is for the non-intervention (control) scenario. The shaded area represents one standard deviation from the mean value of all simulated plots.

**Figure 6.**
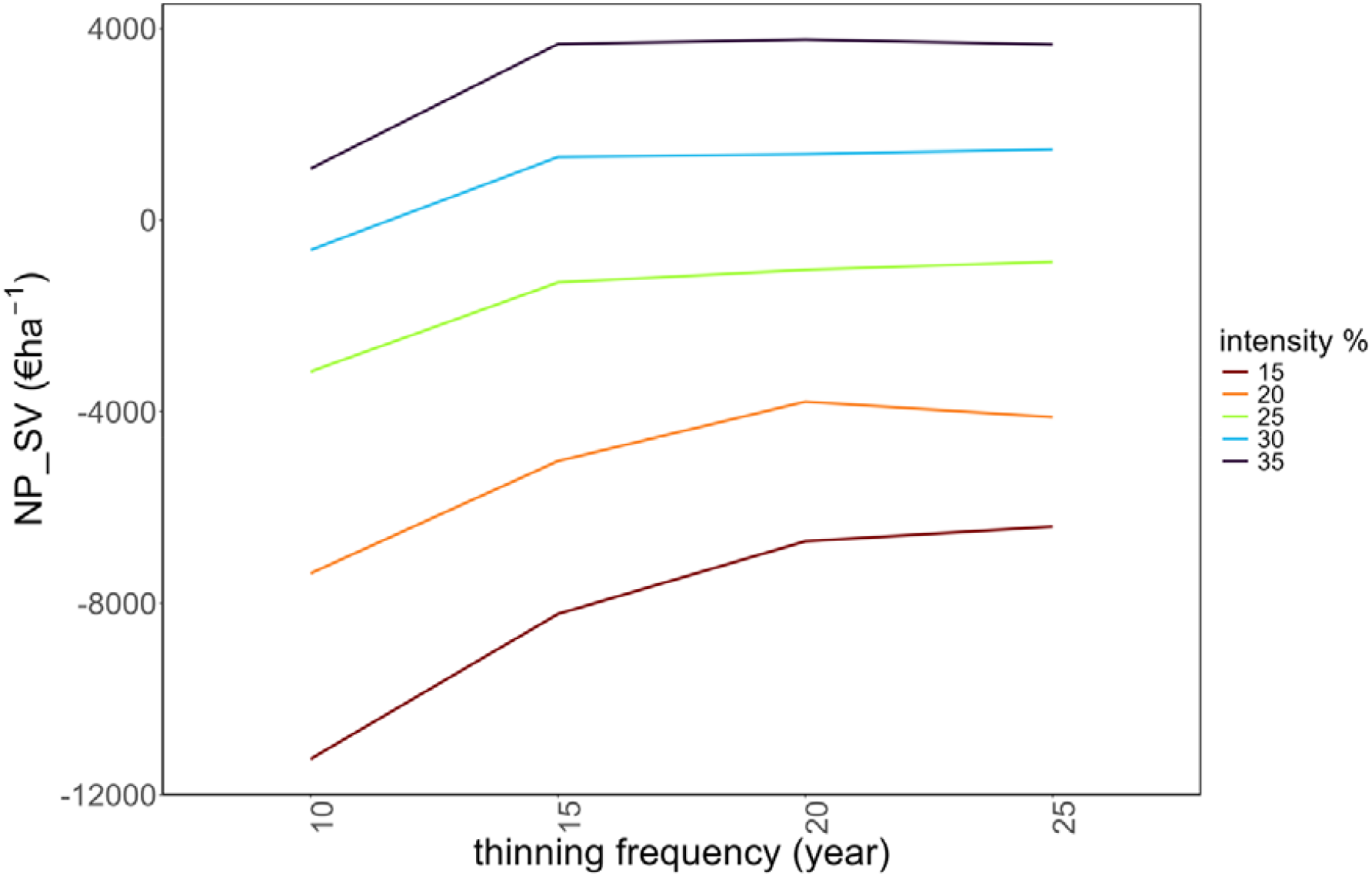
Net Present Stumpage Value (NP_SV) as a function of thinning intensity (removal of 15, 20, 25, 30 and 35% of basal area) and frequency (every 10, 15, 20 and 25 years). Example based on the Monte Amiata plantation.

The average net present values across the various thinning regimes for the case studies of Monte Amiata, Rincine, and Varco San Mauro were as follows: (i) productive function: -2,178 € ha□^1^, 1,777 € ha□^1^, and -431 € ha□^1^, respectively; (ii) aesthetic function: 24,917 € ha□^1^, 28,199 € ha□^1^, and 27,567 € ha□^1^; (iii) erosion protection: 141 € ha□^1^, 215 € ha□^1^, and 163 € ha□^1^; (iv) carbon storage: 28,843 € ha□^1^, 36,743 € ha□^1^, and 35,138 € ha□^1^. The Total Economic Value (TEV), averaging all ES, was therefore 51,723 € ha□^1^, 66,934 € ha□^1^, and 62,438 € ha□^1^ for Monte Amiata, Rincine, and Varco San Mauro, respectively.

The H/D ratio improved as the thinning intensity increased and the frequency decreased, with average values of 63, 66, and 66 for Monte Amiata, Rincine, and Varco San Mauro, respectively. Avoided damages, depending on the thinning regime, ranged between 1,037 € ha□^1^ and 7,031 € ha□^1^ for Monte Amiata (2–18% of TEV), 207 € ha□^1^ and 3,948 € ha□^1^ for Rincine (0.3–7% of TEV), and 570 € ha□^1^ and 8,846 € ha□^1^ for Varco San Mauro (1–16% of TEV).

The financial analysis for individual silvicultural interventions identified the minimum harvested wood volume required to achieve a positive stumpage value in each thinning regime. For example, in the case of the plantation in Varco San Mauro (Table 4), the minimum harvesting volume required for financial efficiency was approximately 160 m^3^ ha□^1^. Yield increases with stand age, reaching around 650 m^3^ ha□^1^ in 60-year-old plantations (Castellani, 1982), as found in the control plots in Varco San Mauro study area.

**Table 4.** Harvested wood volume per thinning intervention (m^3^ ha□^1^) as a function of the intervention year, thinning frequency (every 10, 15, 20 and 25 years), and thinning intensity (removal of 15, 20, 25, 30 and 35% of basal area). Values shaded in grey indicate thinnings with a negative stumpage value. Example based on the Varco San Mauro plantation.

| *Frequency: 10 years* |  |  |  |  |  |
| --- | --- | --- | --- | --- | --- |
| *Thinning year* | *Intensity: 15%* | *Intensity: 20%* | *Intensity: 25%* | *Intensity: 30%* | *Intensity: 35%* |
| n | 59 | 78 | 98 | 118 | 137 |
| n+10 | 76 | 98 | 118 | 135 | 150 |
| n+20 | 95 | 121 | 143 | 161 | 176 |
| n+30 | 115 | 144 | 170 | 191 | 208 |
| n+40 | 132 | 166 | 195 | 219 | 237 |
| n+50 | 146 | 185 | 217 | 244 | 258 |
| n+60 | 160 | 205 | 242 | 262 | 291 |
| n+70 | 171 | 223 | 254 | 293 | 313 |
| n+80 | 179 | 235 | 279 | 305 | 319 |

| *Frequency: 15 years* |  |  |  |  |  |
| --- | --- | --- | --- | --- | --- |
| *Thinning year* | *Intensity: 15%* | *Intensity: 20%* | *Intensity: 25%* | *Intensity: 30%* | *Intensity: 35%* |
| n | 59 | 78 | 98 | 118 | 137 |
| n+15 | 91 | 117 | 143 | 166 | 187 |
| n+30 | 117 | 157 | 192 | 221 | 249 |
| n+45 | 136 | 186 | 233 | 270 | 301 |
| n+60 | 156 | 212 | 275 | 318 | 357 |
| n+75 | 170 | 231 | 298 | 355 | 381 |

| *Frequency: 20 years* |  |  |  |  |  |
| --- | --- | --- | --- | --- | --- |
| *Thinning year* | *Intensity: 15%* | *Intensity: 20%* | *Intensity: 25%* | *Intensity: 30%* | *Intensity: 35%* |
| n | 59 | 78 | 98 | 118 | 137 |
| n+20 | 100 | 134 | 167 | 196 | 223 |
| n+40 | 130 | 176 | 225 | 274 | 317 |
| n+60 | 155 | 210 | 268 | 329 | 394 |
| n+80 | 175 | 236 | 299 | 366 | 433 |

| *Frequency: 25 years* |  |  |  |  |  |
| --- | --- | --- | --- | --- | --- |
| *Thinning year* | *Intensity: 15%* | *Intensity: 20%* | *Intensity: 25%* | *Intensity: 30%* | *Intensity: 35%* |
| n | 59 | 78 | 98 | 118 | 137 |
| n+25 | 109 | 146 | 183 | 222 | 258 |
| n+50 | 141 | 191 | 243 | 296 | 355 |
| n+75 | 170 | 228 | 289 | 352 | 418 |

## 4. Discussion

Effective forest policy and programming requires integrating ES provision into management frameworks (Fares et al., 2015). Our study explored the effects of the intensity and frequency of thinning from below on ES provision in European black pine plantations established in Italy for environmental improvement. It is well-known that thinning from below is one of the most used and effective early-practice in artificial forest stands, especially in conifer plantations to regulate stand density, controlling competition and biomass allocation among trees (Jonard et al., 2006; Cantiani et al., 2015; Cabon et al., 2018).

The 3D-CMCC-FEM model used to simulate stand growth under different thinning regimes has effectively captured the effects of silvicultural interventions on stand structure and dynamics, aligning with findings from previous studies (Dalmonech et al., 2022; Testolin et al., 2023; Saponaro et al., 2025). Diameter growth was positively influenced by heavy-intensity thinning regimes applied at short intervals, reaching the greatest increment with a frequency of 10 years and 35% of basal area removal, while low-intensity regimes proved to be most effective in gaining higher values of stand basal area and AGC (Marchi et al., 2018; Simon and Ameztegui, 2023), also compared to the control scenario (non-intervention). Similar results were obtained by Ameztegui et al. (2017) and del Río et al. (2008), who found that low-intensity thinning did not decrease the final basal area significantly with respect to a non-intervention scenario. Corona (2024) also observed that for optimizing tree density in relation to wood volume growth, a low-intensity regime with relatively frequent thinning interventions (approximately every 10 years) proves to be the most effective. This approach involves moderate biomass removals by each intervention, typically around 15-25% of the stand basal area. In addition, thinning interventions have the benefit of decreasing the competition among trees by reducing the stand density and increasing the stand resistance to extreme events, such as windstorms, potentially leading to a longer-lived forest which represents a long-term biomass pool (Dalmonech et al., 2022; Testolin et al., 2023; Vangi et al., 2024a).

The modeling analyses conducted to economically quantify ES, such as timber production, aesthetic value, erosion control, and atmospheric carbon sequestration, have provided valuable insights for managing European black pine plantations taking into account their multifunctional role. The economic value of provisioning services, particularly timber production, increases with greater basal area removals and longer thinning intervals. A net financial gain is achieved when basal area removal reaches approximately 25% with a 25-year thinning interval, highlighting the higher financial efficiency of more intensive interventions. Thresholds for harvested timber volume that yield a net financial gain are found to be around 160 m^3^ ha^-1^. These relatively high thresholds are due to the significant costs associated with harvesting operations and the need for economies of scale.

Non-provisioning ES (erosion protection, carbon sequestration, and aesthetic/recreational value) tend to decrease with increased basal area removal but benefit from longer intervals between thinning. For the studied stands, the economic values of aesthetic appeal and carbon sequestration far exceed those of timber production and erosion protection, regardless of thinning regime. Specifically, the updated values for aesthetic and carbon sequestration services are, on average, 15-20 times higher than the updated net financial gain of timber production, highlighting the limited trade-offs among these ES in monetary terms. Other studies in the Mediterranean area have found that timber production accounts for only a small part of the Total Economic Value of forests compared with other benefits such as carbon sequestration, watershed protection, and recreation (Croitoru, 2007; Bottalico et al., 2016). The relatively modest monetary value of erosion protection is linked to two factors: (1) the good forest cover in the initial year and its exponential improvement following the first thinning intervention, and (2) the limited social cost associated with erosion compared to other hydrogeological risk regulation functions, such as flood and landslide prevention (Grilli et al., 2020). The Total Economic Value (TEV) of the European black pine plantations in this study for the considered ES is approximately €60,000 per hectare, based on cash flow from 2016 to 2100.

Mechanical stability, assessed by the H/D ratio, improves post-thinning, regardless of the treatment type, although this effect diminishes over time. All thinning treatments improve height-to-diameter ratios and enhance stand stability, reducing potential damages in a range from €207 to €8,846 per hectare. Generally, avoided damage is offset by the TEV of these stands, even when the financial outcome of thinning operations (stumpage value) is negative. This holds true for basal area removal levels of at least 25% in pine plantations. Similar results were announced by Marchi et al. (2018) in the Monte Amiata area, where they found that higher volume and basal area removal lead to higher tree stability and carbon sequestration potential, especially if is considered the carbon fixed in thinned trees. Their results were preliminary due to the short observation period; however, our simulation study seems to confirm their conclusion and align with the literature on the same species and environment (Ruiz-Penado et al., 2013; Marchi et al., 2018, Simon and Ameztegui, 2023).

The analysis of forest ES and their relationships with forest management is a complex task, especially when multiple drivers of change and a wide range of ES are considered for a more complete assessment of forest functions (Nocentini et al., 2022). In our study we focused on the relationship between forest management and ES to avoid confounding effects due to other drivers of change, such as climate change and fires (Morán-Ordóñez et al., 2019). We did not account for potential species change or migration as the simulated period was relatively short for such dynamics, and the study was designed specifically to explore the effects of thinning on the provision of ES and their economic values provided by European black pine plantations. Although the ES we considered represent only a portion of the wide spectrum of benefits provided by forests, we focused on some ES which were less investigated in previous studies, such as soil protection and the aesthetic value of the forest (Marchi et al., 2018; Nocentini et al., 2022; Simon and Ameztegui, 2023).

Finally, the proposed modeling approach economically evaluates multiple ES by integrating standard parameters and coefficients which are specific to the plantations in our case study. Its modular flexibility allows for the inclusion of other drivers of change and additional ES in future applications, such as for example fire hazard reduction, facilitates detailed assessments, and supports sensitivity analyses on variables such as harvesting costs, average assortment prices, interest rates, annual visitor numbers, sediment removal costs, and CO_2_eq prices. Future research could further explore trade-offs among ES in greater detail and under different future climate scenarios.

## 5. Conclusions

This study demonstrates that, in a pine plantation managed for multiple uses, the long-term planning of thinning operations can significantly impact the achievement of management goals. Using a modeling approach, we found that the intensity and frequency of thinning interventions have a substantial effect on the overall production of ES in European black pine stands established through afforestation and reforestation for environmental improvement. Therefore, strategic, long-term planning of thinning operations is essential to ensure a balanced trade-off between wood production and other ES, while maintaining the cost-effectiveness of operations. This includes better monitoring of ES supply and demand, enhanced policy integration, and the development of payment schemes for ES.

The proposed modeling framework serves as an efficient tool for addressing questions that would otherwise require long-term field surveys. By integrating ES provision under various thinning regimes, this approach generates valuable insights to inform the silvicultural tending of stands established through afforestation and reforestation and provides essential information to guide management decisions. While such decisions must be adapted to the unique characteristics of each forest stand, as well as the specific needs and demands of local markets and communities, the framework ultimately offers guidelines for forest planners and managers to ensure the sustained provision of multiple ES.

## CRediT authorship contribution statement

**Elia Vangi:** Writing – review & editing, Writing – original draft, Visualization, Validation, Methodology, Investigation, Formal analysis, Data curation. **Sandro Sacchelli**: Writing – review & editing, Writing – original draft, Methodology, Investigation, Formal analysis, Data curation. **Susanna Nocentini:** Writing – review & editing. **Manuela Plutino:** Writing – review & editing. **Daniela Dalmonech**: Writing – review & editing. **Alessio Collalti:** Writing – review & editing. **Davide Travaglini:** Writing – review & editing, Writing – original draft, Conceptualization. **Piermaria Corona:** Writing – review & editing, Supervision, Resources, Project administration, Investigation, Funding acquisition, Conceptualization.

## Data availability

Data will be made available on request. To the corresponding author.

## Acknowledgments

E.V. and A.C. has been partially supported by MIUR Project (PRIN 2020) “Multi-scale observations to predict Forest response to pollution and climate change” (MULTIFOR, project number: 2020E52THS). A.C. acknowledge also funding by the project OptForEU Horizon Europe research and innovation programme under grant agreement No. 101060554. D.D. and A.C. also acknowledge the project funded under the National Recovery and Resilience Plan (NRRP), Mission 4 Component 2 Investment 1.4 - Call for tender No. 3138 of December 16, 2021, rectified by Decree n.3175 of December 18, 2021 of Italian Ministry of University and Research funded by the European Union – NextGenerationEU under award Number: Project code CN_00000033, Concession Decree No. 1034 of June 17, 2022 adopted by the Italian Ministry of University and Research, CUP B83C22002930006, Project title “National Biodiversity Future Centre - NBFC”. The 3D- CMCC-FEM model code is publicly available and can be found on the GitHub platform at: https://github.com/Forest-Modelling-Lab/3D-CMCC-FEM. The technical support provided by Vincenzo Bernardini, Fabrizio Ferretti, and Chiara Lisa must also be acknowledged.

